# Development, validation, and application of the ribosome separation and reconstitution system for protein translation in vitro

**DOI:** 10.1101/2021.05.31.446494

**Authors:** Brandon M. Trainor, Dimitri G. Pestov, Natalia Shcherbik

## Abstract

The conventional view regarding regulation of gene expression is based on transcription control. However, a growing number of recent studies has revealed the important additional impact of translational regulation. Eukaryotic translational machinery appears to be capable of reprogramming mRNA translation to generate proteins required to maintain a healthy cellular proteostasis under particular physiological conditions or to adapt to stress. Although the mechanisms of such remarkable regulation are beginning to emerge, recent studies have identified the ribosome as one of the major constituents of translation-dependent control of gene expression that is especially important during stress. Built of RNA and proteins, ribosomes are susceptible to environmental and intracellular stresses. How stress-modified ribosomes regulate translation and whether they play a role in stress-induced gene expression remain largely elusive. This knowledge gap is likely due to the lack of an appropriate experimental system. Canonical approaches based on exposing cells or cell-free extracts to stressors provide inconclusive results due to off-target effects of modifying agents. Here we describe a robust and simple in vitro assay that allows separation of yeast ribosomes from other translational machinery constituents, followed by reconstitution of the translation reaction. This **<u>r</u>**ibosome **<u>s</u>**eparation and **<u>r</u>**econstitution assay (RSR) is highly advantageous, as it allows modification of ribosomes without compromising other key translational components, followed by supplementing the ribosomes back into translation reactions containing undamaged, translationally-competent yeast lysate. Besides addressing the impact of ribosome-derived stress on translation, RSR can also be used to characterize mutated ribosomes and ribosomes devoid of associated factors.

## INTRODUCTION

Cell-free translation systems are powerful experimental assets with a wide variety of applications. They allow protein production in a tightly controlled environment using either endogenous transcripts or mRNA reporters. The generated proteins can then be used in subsequent applications like pull-down assays, or analyzed as read-outs of translation reactions addressing roles of cis- and trans-acting factors in translation (Carlson et al. 2012; Chong 2014). Additionally, the cell-free translation reaction allows subsequent supplementation with carefully designed additional factors. This approach is instrumental to identifying the order of molecular events in complex co-translational mechanisms (Shao and Hegde 2014)(Kuroha et al. 2018).

Despite all the advantages of cell-free translation systems, they remain insufficient in dissecting the effects of stress on translational executors. In fact, under stressful conditions, various molecules of the translational machinery undergo modifications (Simms et al. 2014; Tanaka et al. 2007); (Wu et al. 2018);(Yan et al. 2019) (Gu et al. 2014; Endres et al. 2015; Chan et al. 2012). Thus, it is impossible to distinguish between a stressor’s impact on a particular molecule of interest and on other translationally essential elements.

Built of RNAs and proteins, ribosomes are, unsurprisingly, highly susceptible to chemical modifications. Indeed, ribosomes undergo significant modifications when exposed to chemical compounds, metals, or reactive oxygen species (ROS) (reviewed in Shcherbik and Pestov 2019). In addition, in response to a variety of stress conditions, ribosomes and ribosome-bound nascent chains are subject to post-translational protein modifications, such as ubiquitination (reviewed in Dougherty et al. 2020). How performance of *modified ribosomes* is altered during protein synthesis remains largely unknown. This shortcoming is primarily due to the unavailability of a suitable experimental platform that would allow modification of ribosomes *exclusively* and keep other translationally essential molecules intact.

To overcome this technical limitation, we sought to develop a method for purifying ribosomes without affecting their translational activity and placing them back into translationally active ribosome-free yeast lysate charged with an mRNA reporter (schematics in Fig. 1). This approach allows incorporation of the ribosomal modifications into the procedure, followed by precipitating ribosomes. The success of this approach depends on the purification of intact and translationally functional ribosomes from the cell-free extract (CFE).

**FIGURE 1.**
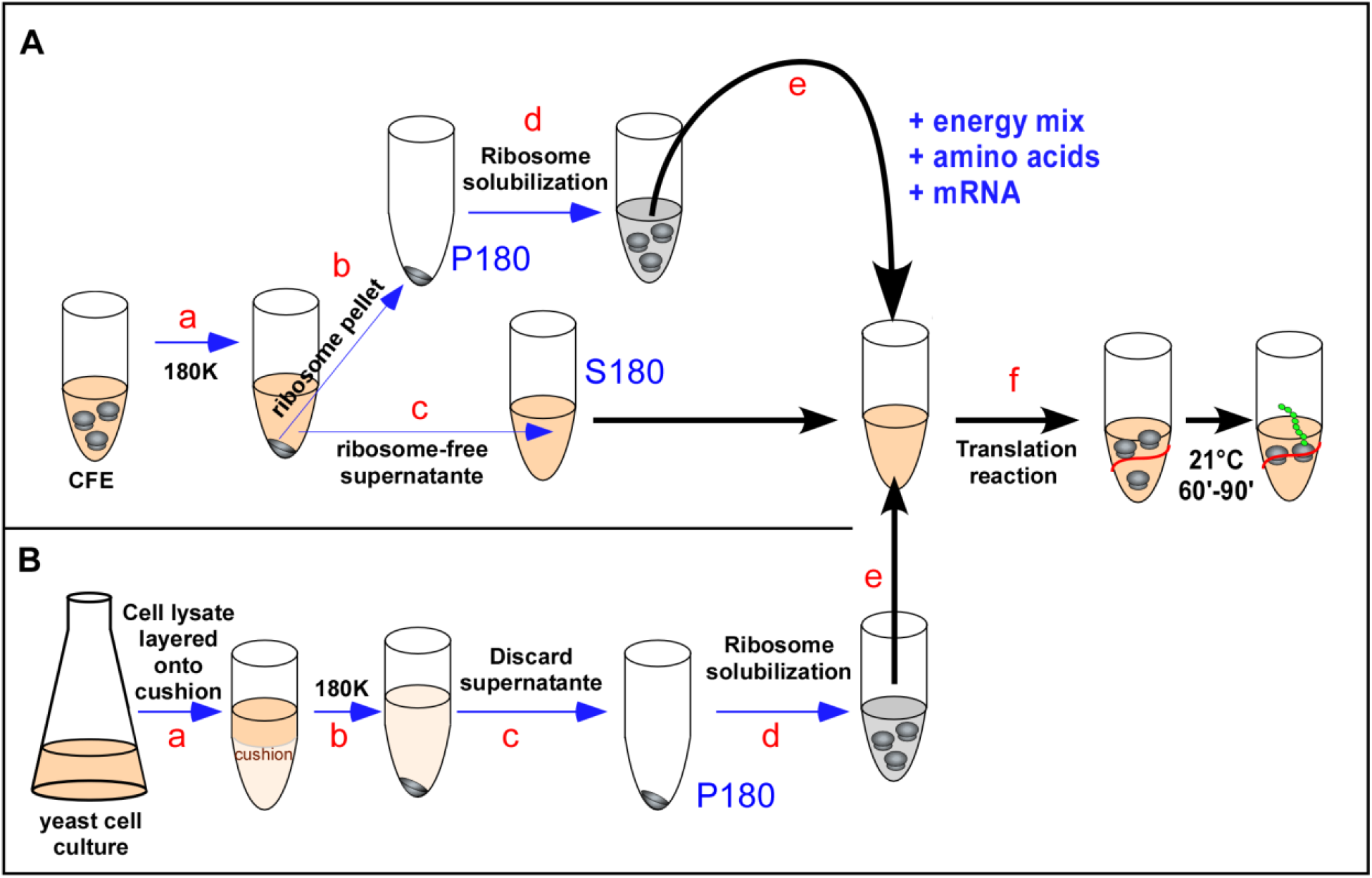
Ribosome separation and reconstitution assay (RSR). (*A*) Ribosomes derived from cell-free extract (CFE) were precipitated by ultracentrifugation at 180,000 × g (180K) for 2 h at 4°C (a), resulting in production of a ribosomal pellet P180 (b) and ribosome-free supernatant S180 (c). Ribosomal pellet P180 was solubilized (d) and added back to S180, along with energy regeneration system, amino acids, and mRNA (e). Translation reactions were carried out at 21°C for 60-90 mins (f). (*B*) Yeast cells were grown and lysed, and total cellular lysate was layered onto 20% glycerol cushion (a). Ribosomes were precipitated by ultracentrifugation at 180,000 × g (180K) for 2 h at 4°C (b). The supernatant was discarded (c), and ribosomal pellet (P180) was solubilized (d) and placed into translation reaction as in (*A*).

The methodology for isolating ribosomes and ribosomal complexes has been described for different organisms and is fine-tuned to each experiment’s goals, which does not always require translational activity of the isolate. Thus, the particular purpose of the isolation dictates the stringency of the ribosome purification protocol. In general, there are two primary strategies to isolate ribosomes, centrifugation and immunoprecipitation (IP), both of which have substantial limitations. Centrifugation-based technology, described by different laboratories, requires lengthy and often numerous spins and thus subjects ribosomes to prolonged exposure to ribonucleases and proteases present in crude cellular lysates. Another limitation of centrifugation-based ribosome isolation is poor solubilization of the resulting ribosomal pellet (Munoz et al. 2017).

On the contrary, the IP approach is fast and avoids pelleting, resulting in soluble ribosomal yields (Oeffinger et al. 2007). However, it demands incorporating a tag on the surface-exposed r-protein, which may interfere with a researcher’s goals. In addition, IP method requires high salt concentrations in the precipitation and elution buffers to avoid pulling down non-specific molecules, implying undesirable stripping of ribosomal co-factors that may perform auxiliary roles during translation (Mazaré et al. 2020; Simsek et al. 2017; Shi et al. 2017).

Here, we report an experimental protocol for purifying translationally competent ribosomes from *Saccharomyces cerevisiae* cells or CFE by one-step centrifugation. Purified ribosomes supplied back into translationally active ribosome-free lysates retained their translational competency and synthetized proteins from various mRNA reporters or endogenous transcripts present in CFE. We term this method the ribosome separation and reconstitution assay (RSR). RSR is simple, rapid, and cost-effective. More importantly, RSR allowed purification of ribosomes exposed to stress. Using the ROS inducer menadione and chemotherapeutic drugs cisplatin as two rRNA modifiers with different characteristics, we demonstrate here the capabilities of the RSR assay in dissecting effects of environmental and intracellular stresses to which ribosomes may be subjected. We also discuss a “ mix-and-match” variation of the RSR assay that may prove instrumental in unraveling the functionality of heterogenous ribosomal pools.

## RESULTS

### Establishment and validation of the RSR

In our previous work, we have developed, validated, and applied a cell-free translation system (CFE) that represents a translationally competent yeast lysate capable of protein synthesis from an mRNA reporter or endogenous cellular transcripts (Trainor et al. 2021). Although this system allows us to address roles of cis and trans factors during protein synthesis, as exemplified in (Trainor et al. 2021), manipulating ribosomes exclusively to research effects of ribosome-directed stress on translation remains elusive.

To develop a system that allows ribosome separation followed by reconstitution of translation in vitro, we used CFE as a starting platform. We aimed to purify ribosomes from CFE by one-step ultracentrifugation and assess their quality and activity during protein synthesis in vitro by returning them into translationally active ribosome-free yeast lysate charged with an mRNA reporter (schematics in Fig. 1A).

#### Preparation of ribosomes purified from CFE by one-step ultracentrifugation

To pellet ribosomes, we centrifuged one aliquot of CFE (∼560 μg of RNA, 1590 μg of proteins) at 180,000 × g for 2 h at 4°C (Fig. 1A-a) in the TLA55 Beckman rotor and collected the supernatant (S180, Fig. 1A-b) and pellet (P180, Fig. 1A-c) fractions. Pelleted ribosomes were solubilized in translation reaction buffer A (20 mM Hepes–KOH, pH 7.4; 100 mM KOAc; 2 mM Mg(OAc)_2_; and 2 mM DTT) (Wu and Sachs 2014) (Fig. 1A-d and see below) and analyzed by northern blotting, along with S180 and complete CFE used as controls. Hybridization with probes specific to 25S and 18S rRNAs verified that the two fractions generated by this ultracentrifugation (180,000 × g for 2 h; 180K-centrifugation hereafter) represent the ribosome-free supernatant (S, Fig. 2A, lane 2) and ribosome-enriched pellet (P, Fig. 2A, lane 3). rRNAs derived from the pellet fraction revealed no signs of degradation (Fig. 2A), suggesting that 2 h of centrifugation does not affect rRNA integrity.

**FIGURE 2.**
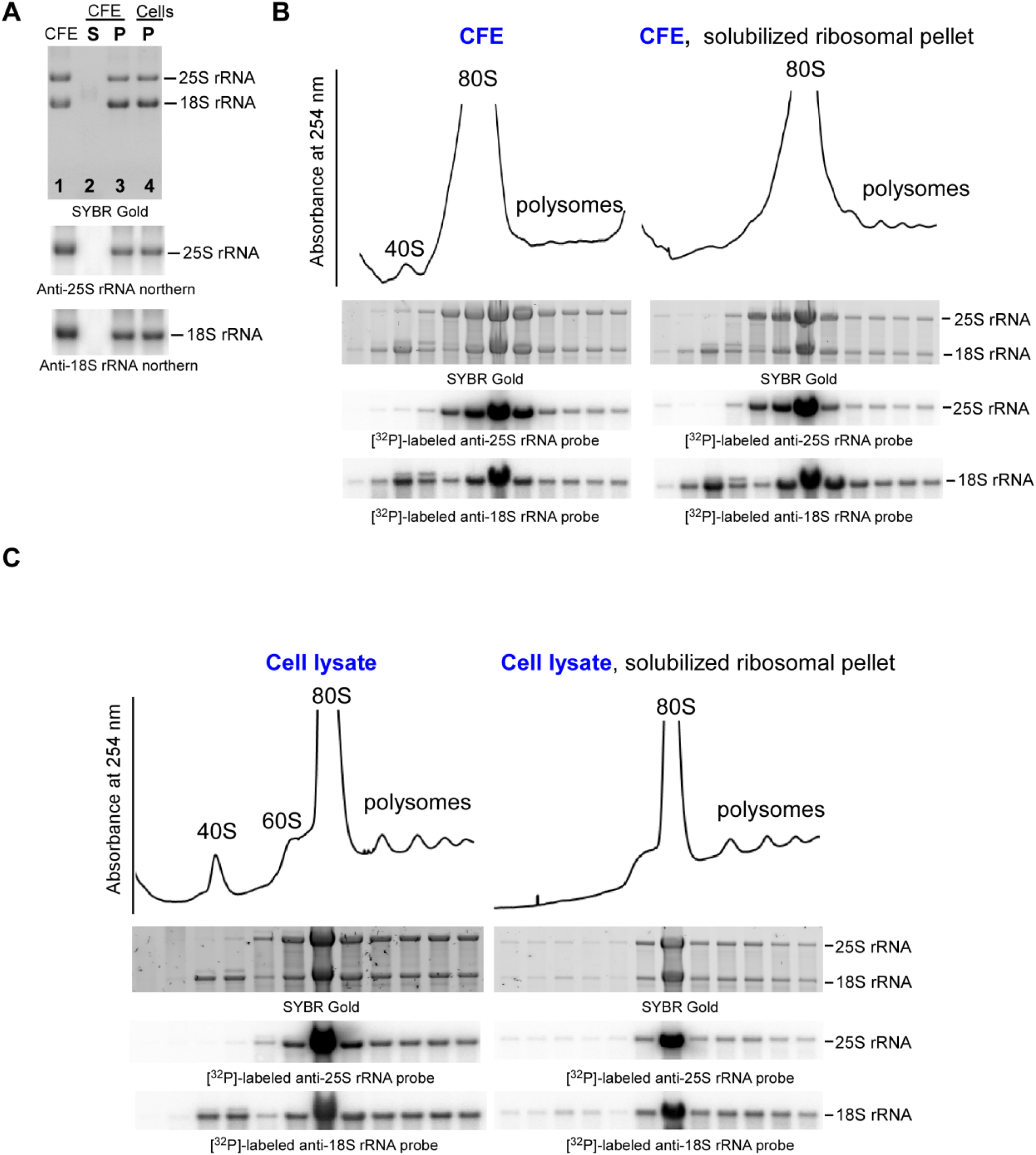
(*A*) One-step ultracentrifugation generates stable ribosomes and ribosome-free lysate. Aliquots of complete CFE (CFE), separated S180 and P180 from Fig. 1A (CFE S P), and just P180 from Fig. 1B (P) were resolved on denaturing agarose gels and analyzed by northern hybridization with indicated probes. Prior to transfer onto nylon membrane, the gel was stained with SYBR Gold. (*B-C*) Ribosomes precipitated by one-step ultracentrifugation exist as 80S monosomes. (*B*) Complete CFE (CFE) and solubilized P180 from Fig. 1A (CFE, solubilized ribosomal pellet) were centrifuged through 15–45% (w/v) sucrose gradients and fractionated with continuous absorbance measurement at 254 nm to visualize ribosomal peaks. RNA was extracted from individual fractions and analyzed by northern hybridization as described in A. (*C*) Sucrose gradient centrifugation analysis performed with the total cellular lysate (cell lysate) and P180 from Fig. 1B (cell lysate, solubilized ribosomal pellet).

#### Characterization of pelleted ribosomes by sucrose gradient centrifugation analysis

Next, using sucrose gradient centrifugation analysis, we examined which ribosomal species were precipitated during 180K-centrifugation (Fig. 1A-a). As a control, we used a CFE sample that was not subjected to ribosome pelleting. Gradients were fractionated into 12 fractions, and RNA was extracted from each fraction and analyzed by northern hybridization with probes specific to 25S and 18S rRNAs. Gradient analysis showed that ribosomes predominantly accumulated in the 80S fraction, with only residual amounts present on polysomes in CFE before and after 180K-centrifugation (Fig 2B). These data demonstrate that 180K-centrifugation is sufficient to pellet polysome-free 80S ribosomes.

#### Optimizing solubilization of pelleted ribosomes

Next we addressed the solubility of pelleted ribosomes, as this technical step was identified as a main limitation of the centrifugation-based ribosome purification approach (Munoz et al. 2017). In fact, we routinely observe that pelleted ribosomes are sticky and difficult to resuspend by pipetting. Due to lack of published detailed information on the resuspension procedure for pelleted ribosomes, we tested the effects of temperature and automated agitation in facilitating ribosomal pellet solubilization. Three CFE-derived ribosomal pellets were incubated in 100 μL of translation buffer A for 30 min at 8°C, 21°C, and 37°C with 1,200 rpm shaking in Eppendorf thermomixers. Subsequent centrifugation of ribosomal solutions at 21,000 × g for 15 min at 4°C did not produce any visible pelleted insoluble debris. The RNA concentrations were similar in all ribosomal suspensions; pellets resuspended at 8°C yielded 5.5 μg/μL of RNA, pellets resuspended at 21°C yielded 5.6 μg/μL of RNA, and pellets resuspended at 37°C yielded 5.2 μg/μL of RNA. Quantifying 18S and 25S rRNAs detected by northern hybridization (Fig. 3A) likewise demonstrated similar levels of these rRNAs regardless of the temperature used during solubilization (Fig. 3B). Northern blot analysis also revealed that ribosomes incubated at 8°C, 21°C, and 37°C for 30 min contained intact 25S and 18S rRNAs with no signs of degradation (Fig. 3A), arguing that the solubilization step (Fig. 1A-d) does not affect rRNA stability.

**FIGURE 3.**
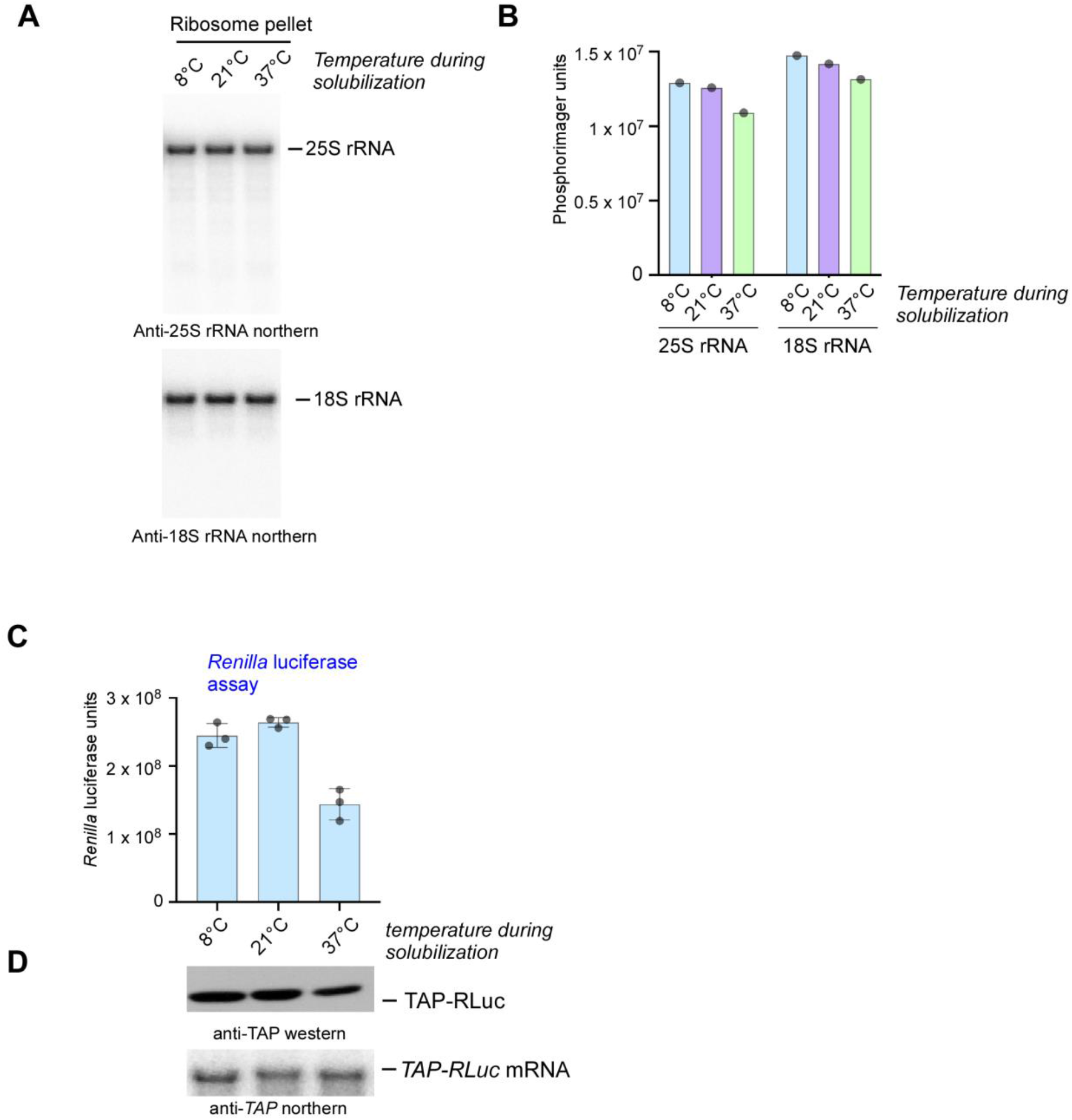
Pelleted and solubilized ribosomes retain their translational competency. (*A*) CFE-derived ribosomal pellets P180 from Fig. 1A were resuspended in translation buffer by shaking for 30 min at the indicated temperatures. Resuspended RNA was analyzed by northern hybridization with 25S rRNA and 18S rRNA specific probes. (*B*) The hybridization signals corresponding to the full-length 25S rRNAs and 18S rRNAs were converted to phosphorimager units and plotted as bar graphs. (*C*) 3 μg of resuspended ribosomes were placed into translation reactions containing ribosome-free translational lysate S180 charged with capped *TAP-RLuc* mRNA (200 ng per reaction). Reaction products were analyzed by the *Renilla* luciferase assay and the data are presented as bar graphs, wherein error bars represent standard error of the mean (SEM) of 3 experiments. (*D*) Proteins and RNA were extracted from the luciferase reactions and further characterized by western blots using anti-TAP antibodies and by northern hybridizations using *TAP*-specific probe.

To examine translational activity of the 180K-pelleted/solubilized ribosomes, we assembled in vitro translation reactions that contained the ribosome-free extract (Fig. 1A-c), 3 μg of ribosomes resuspended at different temperatures, amino acids, and the energy-regeneration system (Fig. 1A-e). Reactions were programmed with 200 ng of *TAP-RLuc* mRNA reporter (*Renilla* luciferase gene fused with TAP-tag). We found that ribosomes solubilized at all temperatures tested synthesized TAP-RLuc as determined by quantitative *Renilla* luciferase assay (Fig. 3C) and confirmed by western blotting (Fig. 3D). However, incubation at 37°C reduced translational activity of ribosomes ∼two-fold (Fig. 3C). The reduced translation activity of 37°C-resuspended ribosomes could be explained by irreversible modifications that likely occur at 37°C or by disassociation of key translation factors (Cox et al. 1973; Danielsson et al. 2015). This result also correlated with poor performance of ribosomes during CFE-based translation at 37°C, further confirming that this temperature is not optimal for ribosomes extracted from BY4741 under low ionic stringency conditions, such as 100 mM KOAc and 3 mM Mg(OAc)_2_ (Algire et al. 2002; Khatter et al. 2014; Pestova et al. 1998; Wu and Sachs 2014). Based on published literature, this buffer composition appears to be optimal to promote correct folding of rRNAs within the ribosomal structure and provide inter-subunit stability (Khatter et al. 2014).

Interestingly, another study demonstrated that translationally active lysates prepared from yeast background strain GRF-18 resulted in higher protein yield and faster kinetics of protein synthesis in vitro at 37°C than at 25°C (Altmann et al. 1989). Therefore, it seems reasonable to propose that different yeast genetic backgrounds might have individual specific temperature requirements. Unless using BY4741 or its derivatives, researchers will have to determine the optimal temperature for the in vitro translation reaction and examine temperature requirements for the ribosome solubilization step of the RSR procedure as illustrated in Fig. 3C.

The important conclusion drawn from this set of experiments is that precipitation of ribosomes from CFE by one-step centrifugation at 180,000 × g recovers stable and active ribosomes. Returning these ribosomes into ribosome-free supernatant reconstitutes protein translation, indicating that RSR approach is effective.

#### Validation of translational activity of ribosomes purified by 180K-centrifugation

Having established P180-ribosome solubilization and buffer composition requirements, we examined the kinetics of protein synthesis using the RSR approach in comparison to complete CFE. The rationale behind this experiment was to monitor the efficiency of protein synthesis over time to identify points at which synthesis is linear and the point at which synthesis plateaus. This is an important characteristic to identify for the RSR system as it defines the optimal time for reaction duration, wherein reliable changes between two or more experimental conditions can be detected.

CFE-pelleted, 21°C-solubilized ribosomes (9 μg of total RNA from P180) were added to ribosome-free extract (S180) charged with 400 ng of capped *TAP-RLuc* mRNA reporter. In parallel, we assembled a reaction with complete CFE using the same amount of *TAP-RLuc* mRNA. Unlike in experiments illustrated in Fig. 3C, in which we measured TAP-RLuc in 15 μL reactions, we assembled 30 μL reactions for the kinetics experiment. Using the *Renilla* luciferase assay, we analyzed 4 μL of the reaction products every 30 min for 3 h. Kinetic analysis revealed that protein synthesis progressed linearly during the first 30–90 min for both RSR and CFE-assembled translation reactions, suggesting that this is the optimal time for reaction duration if samples require comparison (Fig. 4A). Strikingly, we detected higher yield of TAP-RLuc in the RSR reaction than in the complete CFE reaction (Fig. 4A). Besides ribosomes, CFE contain other RNA species (mRNAs, tRNAs); therefore, it is technically impossible to accurately normalize the amount of ribosomes present in complete CFE by measuring RNA concentrations. Thus, we suspected that increased protein synthesis detected in RSR over CFE is likely due to higher concentration of ribosomes in RSR.

**FIGURE 4.**
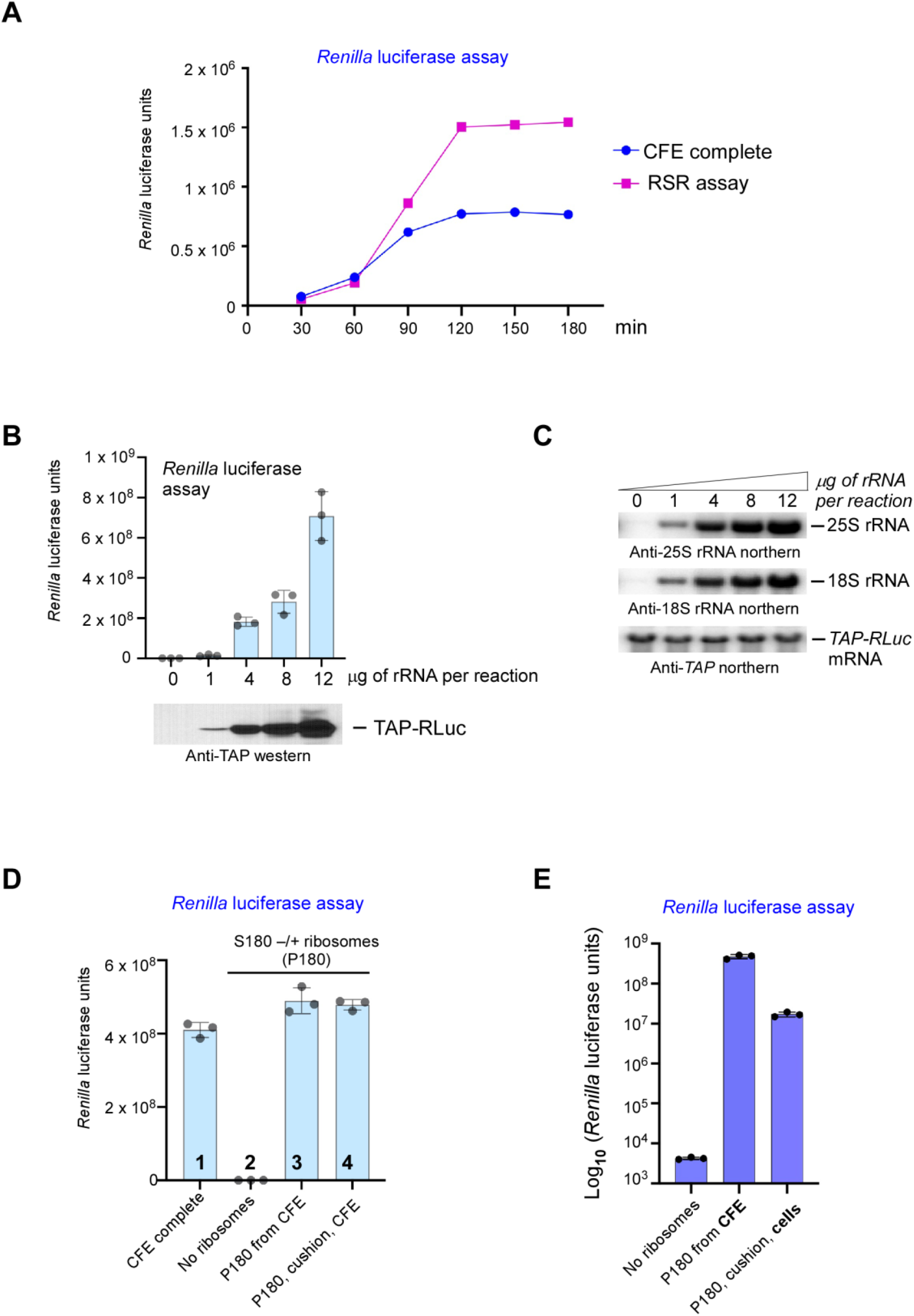
(*A*) Time course of CFE and RSR translation reactions. Translation reactions were assembled with complete CFE (blue) or with ribosomes purified from CFE via 180K-centrifugation (RSR assay, magenta). Both reactions were programmed with 400 ng of capped *TAP-Renilla* mRNA reporter. For the CFE-complete reaction, we used 30 μg of the complete CFE, while the RSR reaction contained 9 μg of purified ribosomes. Reaction aliquots were collected at the indicated time points, and levels of generated reporter proteins were measured by the *Renilla* luciferase assay and plotted as graphs. (*B-C*) Efficiency of translation in RSR reactions depends on the concentration of purified ribosomes. (*B*) Indicated amounts of solubilized ribosomes derived from P180 generated by one-step centrifugation of CFE were added to S180. Each reaction was charged with 200 ng of capped *TAP-RLuc* mRNA reporter. Luciferase signal was measured by the *Renilla* luciferase assay and is presented as bar graphs. Error bars represent SEM of 3 experiments. (*C*) Total RNA was extracted from the RSR/luciferase reactions from (B) and analyzed by northern hybridizations using probes specific to 25S rRNA, 18S rRNA, and *TAP-RLuc* mRNA. (*D*) Centrifugation of ribosomes through glycerol cushion does not affect their translational activity. Translation reactions were assembled with the complete CFE (lane 1) or with 9 μg of purified ribosomes in the RSR format (lanes 2–4). For RSR, ribosomes were pelleted directly from CFE (3) or by centrifugation of CFE through a 20% glycerol cushion (4). In (2), no ribosomes were added into the RSR reaction. Each RSR reaction contained 400 ng of capped *TAP-RLuc* mRNA reporter. Reaction products were analyzed as in (B). (*E*) Ribosomes purified from cells retain their translational activity. RSR translation reactions were assembled with 9 μg of ribosomes pelleted directly from the complete CFE (P180 from CFE) or with 9 μg of ribosomes purified from cell lysate by 180K-centrifugation through 20% glycerol cushion (P180, cushion, ribosomes). In the control RSR reaction, no ribosomes were added. Each RSR reaction contained 400 ng of capped *TAP-RLuc* mRNA reporter. Reaction products were analyzed as in (*B*).

To confirm that translation efficiency depends on ribosome concentration, we assembled RSR reactions with different amounts of ribosomes placed into ribosome-free supernatant S180 changed with 200 ng of capped *TAP-RLuc* mRNA reporter (Fig. 1A-e). We assessed the production of TAP-RLuc using the quantitative *Renilla* luciferase assay and chemiluminescent western blotting. As expected and consistent with the previous experiment (Fig. 2A), neither luminescent signal nor TAP-RLuc protein were detected in the reaction containing S180 only (Fig. 4B, lane1 /bar 1), suggesting that S180 lacks endogenous ribosomes after the 180K-centrifugation procedure. As predicted, reporter synthesis efficiency increased with increasing ribosomal content, and the highest amounts of TAP-RLuc were detected in the reaction containing the highest concentration of ribosomes (i.e., 12 μg; Fig 4B). Northern hybridization of RNA extracted from these RSR reactions confirmed gradual increase of 18S and 25S rRNAs (Fig. 4C, top) and verified the stability of *TAP-RLuc* mRNA post reaction (Fig. 4C, bottom).

Taken together, these data suggest that for most efficient protein synthesis using the RSR format of cell-free translation, multiple components must be carefully optimized, including mRNA concentration, reaction temperature, and reaction time.

#### Translational activity of P180-derived ribosomes after centrifugation of CFE through 20% glycerol cushion

Purifying ribosomes via centrifugation through a cushion is routinely used in various biochemical applications (Mehta et al. 2012; Khatter et al. 2014; Jenner et al. 2012). In fact, like IP, this approach allows separation of ribosomes from low molecular weight molecules, including endogenous and exogenous cellular components, which might interfere with the translation reaction. For example, treating CFE with ribosome-modifying compounds (discussed below) would require removing the drug prior to adding ribosomes to the translation reaction. Thus, the ability to use cushion-centrifuged ribosomes would be a powerful feature of the RSR translation system.

To assess the activity of ribosomes precipitated by ultracentrifugation through a cushion in the RSR reaction, we either directly precipitated ribosomes from CFE as described above or centrifuged CFE through a 20% glycerol cushion at 180,000 × g for 2 h. The resulting P180 pellets were solubilized, and 9 μg of P180 were added to the ribosome-free translational lysate (S180) along with 400 ng of capped *TAP-RLuc* mRNA reporter. *Renilla* luciferase assay revealed that ribosomes precipitated by centrifugation through the cushion were as active as those pelleted directly from CFE (Fig. 4D, bars 3 and 4). Consistent with the kinetics experiment (Fig. 4A), translational products resulting from the RSR reactions were slightly more abundant than those generated in complete CFE (Fig. 4D, bars 1 and 3).

#### Activity of ribosomes purified from yeast cell culture in RSR translation reactions

To establish, validate, and optimize RSR format of cell-free translation reactions, we used translationally active CFE lysate as a starting platform. The next question we wanted to address was whether ribosomes extracted directly from cell culture would be active in RSR format. Total cellular lysates prepared by conventional glass bead-beating lysis technique were centrifuged through a 20% glycerol cushion at 180,000 × g for 2 h at 4°C to separate heavy ribosomal particles from low molecular weight cellular contaminants (Fig. 1B-a). Similarly to CFE-purified ribosomes, cell-derived ribosomes demonstrated no signs of degradation (Fig. 2A, lane 4) and were detected in the 80S fraction of the sucrose gradient (Fig. 2C). Cell-derived ribosomes (9 μg) placed into translation reactions charged with 400 ng of capped *TAP-RLuc* mRNA demonstrated translational competency, as measurable luminescent signal was achieved (Fig. 4E, bar 3). However, we observed a significant decline (∼30 times) in reporter synthesis efficiency with cell-derived ribosomes compared to those isolated from CFE (Fig. 4E). Nevertheless, the *Renilla* luciferase signal was still high, suggesting that using cell-derived ribosomes is a reasonable alternative to CFE-purified ribosomes.

### Practical applications of the RSR assay

The established RSR system (Fig. 1-4) can be used for various applications. For example, the RSR-based approach allows modification of ribosomes with stressor like oxidants or chemotherapeutic drugs before supplying them into translation reactions. To illustrate RSR’s application in assessing activity of modified ribosomes, we tested two RNA modifiers: 1) the cell-permeable drug menadione (vitamin K3, or K3), which promotes oxidation of RNA and proteins in cells (Shedlovskiy et al. 2017b; Smethurst et al. 2020; Zinskie et al. 2018); and 2) cell-impermeable drug cisplatin, also known to modify nucleic acids, including rRNAs (Dedduwa-Mudalige and Chow 2015; Melnikov et al. 2016). Purifying translationally active ribosomes from cell cultures (for menadione treatment) or CFE (for cisplatin treatment) allowed us to analyze both drugs’ effects on ribosome-regulated translation efficiency.

### Reduced translation in ribosomes treated with menadione

Menadione is a pro-oxidant used as an extracellular stressor of yeast cells because of its stability in the medium during yeast culture treatment and high cell wall/membrane permeability (Jamieson 1992). In our experience, menadione has advantages over the commonly used hydrogen peroxide due to its stability in solution during storage, which improves data reproducibility among experiments (Shedlovskiy et al. 2017b; Smethurst et al. 2020; Zinskie et al. 2018). Previous studies have found that treating yeast cultures with high doses of menadione (up to 600 μM) triggers excessive rRNA fragmentation (Mroczek and Kufel 2008; Shedlovskiy et al. 2017b) followed by induction of the apoptotic program (Mroczek and Kufel 2008). On the contrary, treating yeast cultures with low doses of menadione (25–50 μM) does not affect cell viability and is accompanied by 25S rRNA cleavage specific for the expansion segment ES7L (Shedlovskiy et al. 2017b). Moreover, we found that menadione-induced ES7L-cleavage does not inhibit translation (Shedlovskiy et al. 2017b), suggesting that limited oxidation of ribosomes might play a regulatory role during protein synthesis.

Unlike the oxidant hydrogen peroxide, which directly modifies its targets, the redox-cycling agent menadione promotes oxidation indirectly by affecting the primary cellular antioxidant glutathione, resulting in ROS accumulation (Kim et al. 2014; Ochi 1996; Morris et al. 2014). Therefore, menadione-induced ribosome oxidation can occur only in the cellular context, since the whole-cell environment is required to generate ROS that modify ribosomes. Therefore, to study the activity of ribosomes oxidized by a menadione-dependent network in the cell-free setting, we applied the strategy illustrated in Fig. 1B.

#### In vitro [^35^S]-Met/Cys incorporation into nascent polypeptides as a read-out of translation

Exponentially growing yeast cultures were treated with 50 μM and 100 μM menadione for 2 h at 30°C or remained untreated. Cells were lysed by glass bead shearing and lysates were centrifuged through 20% glycerol cushion prepared in translation buffer to precipitate ribosomes. Ribosomal pellets were resuspended in the translation buffer by shaking at 21°C for 30 min (Fig. 3C), and same amounts of RNA (12 μg, Fig. 4B) were added to the ribosome-free translational lysates (Fig. 1B-d). These lysates were generated by ultracentrifuging CFE at 180,000 × g for 2 h as described above (Fig. 1A-c). The translation reactions were also supplied with energy mix and amino acids with radioactively labeled methionine (Met) and cysteine (Cys). Aliquots of the reaction products (4 μL) were collected at 5 min, 30 min, and 45 min post-reaction, proteins were precipitated with TCA and the amounts of labeled nascent polypeptides (trapped on filters) generated from endogenous mRNAs present in CFE were measured by scintillation counting. As expected, amounts of [^35^S]-Met/Cys-labeled polypeptides increased over time when ribosomes isolated from untreated cells were used (Fig. 5A, red curve). Ribosomes extracted from cells treated with 50 μM menadione were also translationally active, generating ∼1.3 times less total protein over time than untreated ribosomes, while ribosomes isolated from cells treated with 100 μM menadione were inactive (Fig. 5A, blue and green curves). These data are consistent with our previous observation that 100 μM menadione treatment promotes rRNA fragmentation and significantly affects cell viability (Shedlovskiy et al. 2017b).

**FIGURE 5.**
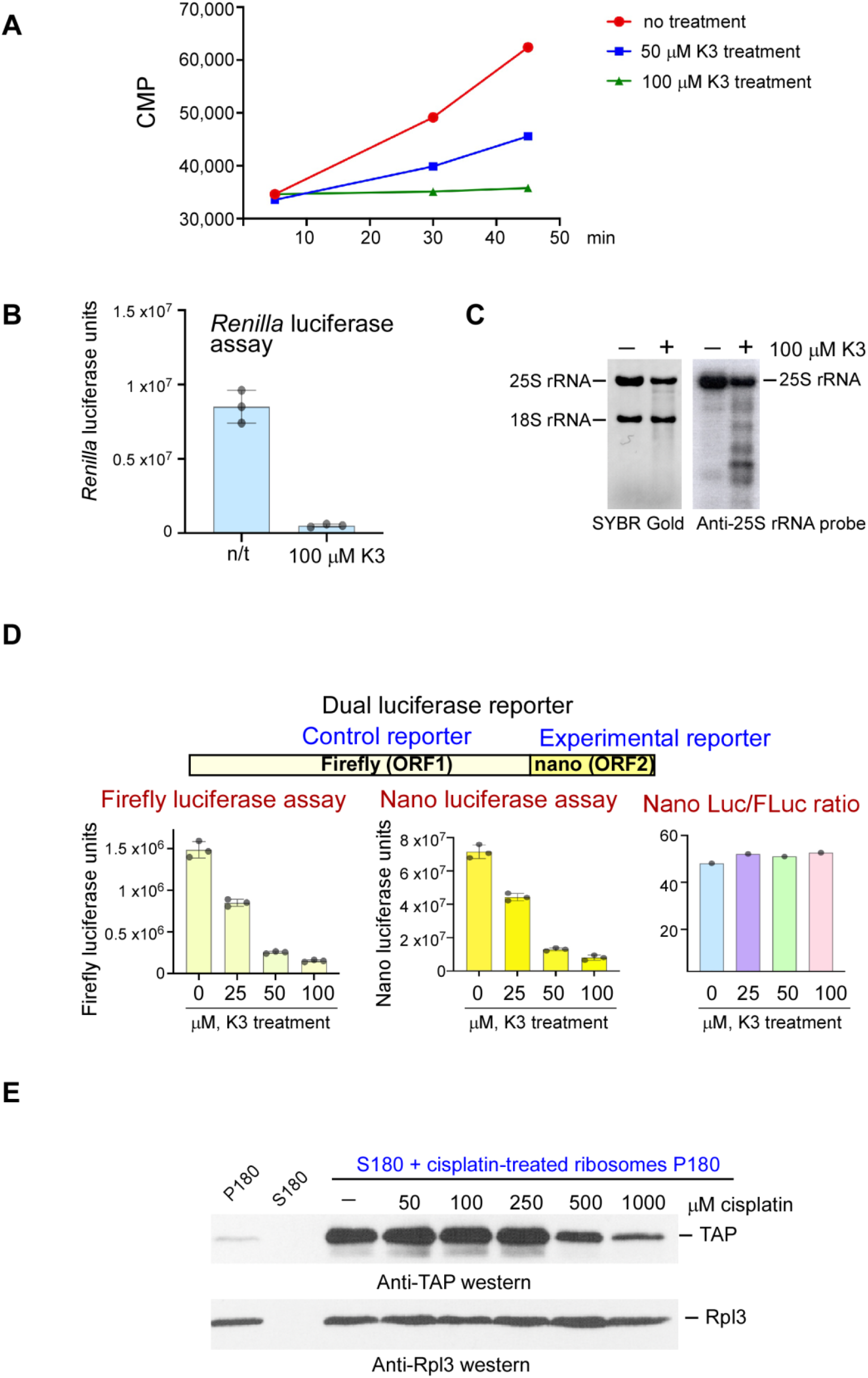
Ribosome translational activity decreases upon treatment with menadione or cisplatin in dose-dependent manner. (*A*) Mid-log wild-type cells BY4741 grown in YPD were treated with 50 μM or 100 μM menadione for 2 h at 30°C or left untreated. Cells were lysed, and ribosomes were precipitated by ultracentrifugation through 20% glycerol cushion as illustrated on Fig. 1B. Ribosomal pellet P180 was resuspended in translation reaction buffer, and 12 μg of ribosomes were added into translationally-active ribosome-free lysate S180 prepared from CFE (Fig. 1A), along with amino acids containing labeled [^35^S]-Met/Cys. Reactions were incubated at 21°C, 4 μL aliquots were taken at indicated time points, and proteins were precipitated by TCA. Incorporation of [^35^S]-Met/Cys into nascent peptides were measured by scintillation counting; CPM (count per minute) values were plotted as graphs. (*B*) Ribosomes were extracted from yeast cultures treated or not (n/t) with 100 μM menadione as described in (A). 9 μg of resuspended ribosomes were added into translation reactions containing S100 and 400 ng of capped-*TAP-RLuc* mRNA reporter. Reaction products were analyzed by the *Renilla* luciferase assay and the data are presented as bar graphs, wherein error bars represent SEM of 3 experiments. (*C*) RNA was extracted from ribosomal suspension from (B) and resolved on a denaturing agarose gel, transferred onto nylon membrane and hybridized with 25S rRNA-specific probe (right). Prior to transfer, gel was stained with SYBR Gold (left). (*D*) *top* Schematics for dual luciferase reporter Firefly-nano luciferase. The control reporter (ORF1) and the experimental reporter (ORF2) are shown. Molecular weight of each reporter is indicated. *bottom* Mid-log wild-type cells BY4741 grown in YPD were treated with 25 μM, 50 μM or 100 μM menadione for 2 h or left untreated (n/t). Ribosomes were extracted, solubilized and 9 μg were placed into translation reaction as described in (*B*). 400 ng of the capped dual firefly-nano luciferase reporter mRNA was added into each reaction. Reaction products were analyzed by Nano-Glo Dual-Luciferase Reporter assay. The data are presented as bar graphs for firefly luciferase (left) and nano luciferase (middle). Error bars represent SEM of 3 experiments. Luminescence signals derived from nano luciferase (experimental reporter) were normalized by luminescence signals derived from firefly luciferase (control reporter) and presented as Nano-Luc/FLuc ratio on the right panel. (*E*) Complete CFE was treated with cisplatin at the concentrations indicated in the figure. Drug-treated CFE was centrifuged through 20% glycerol cushion at 180,000 × g for 2 h, and ribosomal pellets were washed and resuspended in the translation reaction buffer. 9 μg of ribosomes were added to the translationally active ribosome-free lysate S180 charged with 300 ng of capped *TAP* mRNA reporter. Reaction products were analyzed by western blotting with anti-TAP antibodies and control anti-Rpl3 antibodies.

#### mRNA reporters as a convenient tool to assess translation in vitro

Alternatively to [^35^S]-Met/Cys protein labeling approach, the translation reaction can be charged with capped mRNA reporter and ribosomes isolated from stressed or unstressed cells. In this setting, researchers can measure production of a particular protein using the appropriate technique.

Renilla *luciferase reporter*. We supplied 400 ng of m7G-*TAP-RLuc* mRNA and 9 μg of ribosomes extracted from 100 μM menadione-treated or from untreated cells into translation reactions (Fig. 1B-d) and measured the luminescence produced by synthetized TAP-RLuc using the *Renilla* luciferase assay. Consistent with previous data (Fig. 5A), luciferase signal detected in reactions driven by menadione-treated ribosomes was significantly lower than that of ribosomes extracted from untreated cells (Fig. 5B). Northern blot analysis of RNA extracted from translation/luciferase reactions revealed excessive fragmentation of 25S rRNA in samples containing ribosomes from 100 μM menadione-treated cells (Fig. 5C), consistent with our published results (Shedlovskiy et al. 2017b)

#### Dual firefly-nanoluciferase reporter

We also tested a dual mRNA reporter in the RSR system supplemented with menadione-modified ribosomes. Dual reporters are commonly used during gene expression analysis, as they represent a powerful experimental tool that allows data normalization by calculating the ratio of the experimental reporter (ORF2) over the control reporter (ORF1) (schematics in Fig. 5D). This helps in data interpretation, as it takes into consideration unavoidable experimental variability. In this regard, luciferase-based dual reporters, such as the firefly-nanoluciferase used here, bring additional convenience and simplicity to the procedure, allowing simultaneous measurement of two enzymes within a single sample. In dual assays, the activity of the primary reporter (nanoluciferase) reflects the experimental conditions studied while the activity of the control reporter (firefly luciferase) represents the baseline, providing an essential internal control.

400 ng of capped firefly-nanoluciferase (*FLuc-nanoLuc*) mRNA-reporter was added into ribosome-free translationally active lysate (S180, Fig. 1A-c) along with 9 μg of ribosomes extracted from cells treated with 25 μM, 50 μM, and 100 μM menadione. As a control, we used ribosomes extracted from untreated cells. Reaction products were analyzed by the Dual-Luciferase Reporter assay from Promega. We measured the luminescent signal derived from firefly luciferase (Fig. 5D, left panel), then measured nano-luciferase luminescence (Fig. 5D, middle panel). Synthesis of both reporters correlated negatively with increasing menadione concentrations during treatment (Fig. 6D, left and middle panels). Normalizing nanoLuc to FLuc signals revealed a similar ratio in every reaction, confirming low experimental variability of RSR (Fig. 6D, right panel). Moreover, the RSR system’s capability to synthesize FLuc-nanoLuc (81 kDa), which is larger than *Renilla* luciferase (56 kDa), indicates the robustness of this system.

#### Reduced translational efficiency of ribosomes from CFE with high-dose cisplatin

Cisplatin is a chemotherapeutic drug widely used to treat many types of cancer (Dasari and Tchounwou 2014). Although the main target of cisplatin is DNA, it can also modify RNA, in particular rRNAs (Mezencev 2015).

Here, we examined the effects of cisplatin on the translational ability of ribosomes in the RSR system. Since cisplatin is membrane-impermeable, we treated CFE (instead of cells, as for menadione) with various concentrations of this drug for 2 h at 21°C. Untreated CFE was used as a control. To separate cisplatin-modified ribosomes from other CFE components and excess drug, ribosomes were precipitated by one-step centrifugation through 20% glycerol cushion. Ribosome-enriched pellets were washed and resuspended in 100 μL of translation buffer A with shaking at 21°C for 30 min; the 9 μg of rRNA were placed into cisplatin-untreated ribosome-free lysate S180 (Fig 1A-c). The reactions were charged with 300 ng of capped *TAP* mRNA reporter. Reactions were incubated at 21°C for 90 min (Fig. 1A-f) and reporter protein synthesis was analyzed by western blotting using anti-TAP antibodies, with antibodies that detect Rpl3 used as an internal control. As expected, no protein signal was detected in reactions containing only ribosomes or only S180 (Fig. 5E, lanes 1-2), while addition of ribosomes to S180 resulted in strong TAP-reporter synthesis (Fig. 6E, lane 3). Ribosomes’ translational ability declined only when cisplatin was used at high concentrations (0.5 mM and 1 mM, Fig 6E, lanes 7–8), while treatment with 50 μM, 100 μM, and 250 μM had no visible effects reporter levels (Fig. 6E, lanes 4-6).

Previous studies have identified rRNA sites susceptible to cisplatin modifications. For example, Melnikov and colleagues revealed 2.6 Å-resolution crystal structures of bacterial 70S exposed to cisplatin, which demonstrated the drug’s ability to stably intercalate into rRNA structures (Melnikov et al. 2016). Similarly, Rijal and Chow used in vitro and in vivo experimental systems to show that cisplatin can bind both purified 30S subunits and those that are in the 70S ribosomal complex (Rijal and Chow 2009). Furthermore, in yeast, cisplatin bound to RNA more efficiently than to DNA (Hostetter et al. 2012). Taken together, our data demonstrating reduced ribosomal activity upon exposure to cisplatin in the RSR reactions, along with the growing evidence that cisplatin binds RNA, help explain mechanisms of cisplatin toxicity in cells.

## DISCUSSION

We have devised a unique yeast-based biochemical approach (outlined in Fig. 1) that allows purification of ribosomes from cells or cell-free extracts. These ribosomes can be returned into undamaged, translationally active ribosome-free cell lysates charged with an mRNA reporter or with radioactively labeled amino acids, after which the generated proteins are analyzed. This established “ ribosome separation and reconstitution” format of the translation reaction was termed the RSR assay or RSR reaction.

We demonstrated that ribosomes purified from CFE or cell cultures by one-step centrifugation remain stable (Fig. 2) and retain their translational activity (Figs. 3-5). As such, translation reactions reconstituted in vitro with ribosome-free lysates and pellet-derived ribosomes result in efficient translation of both endogenous CFE-derived transcripts (Fig. 5A) and various mRNA reporters (Figs. 3-5), confirming the efficiency and flexibility of RSR.

Interestingly, an approach similar to ours was undertaken by another group, with rabbit reticulocyte lysate used to generate ribosome-free supernatants containing factors sufficient for translation, and with human cells serving as ribosome donors (Penzo et al. 2016). The mammalian protocol used overall similar conditions to pellet ribosomes, namely, centrifuging at 140,000 × g for 5 h (Penzo et al. 2016). Examining rRNAs and r-protein Rpl3 content as a read-out of ribosomal content in the S180 supernatant generated by centrifuging yeast CFE at 180,000 × g for 2 h revealed that these conditions are sufficient to generate yeast lysate devoid of ribosomes (Figs. 2A, 4B&C). Functional assays supported these data, as no protein reporter products were detected in translation reactions that lack exogenously added ribosomes (Fig. 4B, D, E; 5E).

Unlike ribosomes purified by pelleting from CFE (directly or through glycerol cushion), those extracted from cells via conventional glass bead-beating procedure were approximately an order of magnitude less efficient in the translation reaction (Fig. 4E). This calls for improving the method. Optimizing buffer composition for bead-beating lysis could be a reasonable strategy to recover more active ribosomes from cells. Another possibility is applying a spheroplasting-based cell lysis approach, which requires enzymatic digestion of the cell wall followed by lysis using osmotic pressure, freeze-thawing, or other disrupting strategies (Darling et al. 1969; Mann et al. 1972; Izawa and Unger 2017). Finally, the cryogenic lysis technique that has proven to be a very powerful approach for maintaining translational activity (Trainor et al. 2021) can be used. However, this method requires a large sample volume, is laborious, and is limited by the number of samples that can be processed simultaneously. Thus, there is still room for optimizing in the RSR method.

We tested RSR with ribosomes subjected to oxidative stress or to the chemotherapeutic drug cisplatin. The results presented in Fig. 5 demonstrate that both conditions modified the purified ribosomes, affecting their translational activity in a dose-dependent manner. It will be interesting to identify these induced aberrations to rRNAs or r-proteins. Another question of interest is investigating modified ribosomes’ properties during translation of challenging sequences, such as translating mRNA with stalls, rare codon stretches, and programmed ribosome frameshifting. Dual-luciferase reporters, successfully used in this study (Fig. 5D), will be instrumental in addressing these questions.

To the best of our knowledge, a yeast-based RSR-like protocol has never been reported before. Considering that yeast cells can be cultured in large quantities and are very amenable to genetic manipulations, our protocol provides significant methodological advances to the fields of translation, ribosome biology, and protein quality control.

In conclusion, the RSR assay offers a robust and easy-to-use tool for assessing translational competency of modified or damaged ribosomes without the risk of contaminating other critical factors. Data from our lab demonstrated that this approach is also applicable for assaying translation properties of genetically perturbed ribosomes, such as those that contain mutations/deletion of non-essential r-proteins and ribosome-bound co-factors (B.M.T. and N.S personal observations). Overall, the RSR assay will undoubtedly serve as a powerful tool for dissecting translational properties of modified ribosomes and beyond.

## MATERIALS AND METHODS

### Yeast strain, medium, yeast culture treatment

We used YPD media (1% yeast extract, 2% peptone, and 2% dextrose) that was sterilized by filtration through a 0.2 μc filter system (“ Rapid-Flow” from Thermo Scientific). Wild-type BY4741 (*MAT*a *his3–1 leu2– 0 met15– 0 ura3– 0*) was purchased from Open Biosystems. For experiments with menadione treatment, overnight BY4741 yeast cultures were diluted with fresh YPD at A_600_ of ∼0.3 and grown for an additional 2 – 4 h at 30°C. Various concentrations of menadione (indicated in Figures and Figure legends) were added to the cultures; cells were grown for an additional 2 h at 30°C agitating, harvested, washed with water, and lysed.

### Antibodies, chemicals and enzymes

The following antibodies were used: Peroxidase anti-peroxidase complex (PAP) (Sigma, cat# 1291) to detect TAP; anti-Rpl3 (ScRPL3, Developmental Studies Hybridoma Bank, University of Iowa); anti-mouse IgG-HRP (GE-Healthcare, Cat# NA931).

Cisplatin was purchased from Sigma (PHR1624), menadione from Enzo (ALX-460-007-G010), DTT from Sigma (D0632), SYBR Gold from Thermo Fisher Scientific (S11494); TRI-REAGENT-LS was from Molecular Research Center Inc (cat# TS 120); formamide was purchased from Sigma (cat# 47670-25ML-F) and stored in 1 ml aliquots at -80 °C. EasyTag EXPRESS^35^S Protein Labeling mix, [^35^S]-, 2 mCi, 11 mCi/ml was obtained from Perkin Elmer (cat# NEG772002MC).

RiboLock (cat# EO0381), DreamTaq PCR 2x master mix (cat# K1071), 100 mM ATP solution (cat# R0441), 100 mM GTP solution (cat# R0461), proteinase K (cat# EO0491) and all the restriction enzymes and were obtained from Thermo Fisher Scientific. Creatine phosphokinase was purchased from BioVision, creatine phosphate from VWR (cat# 97061-328). 1 mM solution of 20 Essential amino acids, complete and minus Methionine and Cysteine were from Promega (cat# L4461 and L5511, respectively); Ambion mMESSAGE mMACHINE™ T7 Transcription kit was purchased from Thermo Fisher Scientific (cat# AM1344); Crescendo chemiluminescent HRP detection reagent was from Millipore Sigma (cat# WBLUR0100); DNA clean and concentrator kit was from ZYMO RESEARCH (cat# D4004); PD-10 columns Sephadex G-25 (20×80 mm) were from GE Healthcare (cat# 17085101).

### Plasmids

pYes2 was purchased from Invitrogen. To generate pYes-TAP, we amplified the TAP sequence using the pBS1761 plasmid (a kind gift of Dr. Mike Henry) as a template with the forward primer containing the BamH1 site and the reverse primer containing Xho1 site; PCR TAP-product was cloned into pYes between BamH1 and Xho1. We also amplified the TAP sequence using the pBS1761 plasmid with a forward primer containing BamH1 and the reverse primer containing the Xho1 site. The stop codon was omitted from the TAP sequence. PCR TAPNoStop product was cloned into pYes between BamH1 and Xho1, resulting in pYes-TAPNoStop construct. *Renilla* luciferase gene was amplified by PCR from pJD375 with a forward primer containing Xho1 site and reversed primer containing Xba1 site and cloned into pYes-TAPNoStop construct between Xho1 and Xba1 sites, resulting in pYes-TAP-RLuc fusion. The sequences of pYes-TAP and pYes-TAP-RLuc were verified by sequencing.

To generate the dual-luciferase reporter Firefly Luc – nano Luc, firefly luciferase gene was amplified using pJD375 plasmid as a template (a kind gift of Dr. Jonathan Dinman) with the forward primer containing HindIII site and the reverse primer containing BamH1 site, whereby the reverse primer was designed to anneal upstream of the firefly luciferase gene stop codon. The PCR product was cloned into pYes2 between BamH1 and HindIII, resulting in the pYes-FLucNoSTOP construct. Nano-luciferase was amplified using pF4Ag NanoLuc plasmid from Addgene (cat # 137777) as a template. The forward primer contained the BamH1 site, while the reverse primer contained the Xho1 site. The PCR product was cloned into the pYes-FLucNoSTOP construct between BamH1 and Xho1 sites. The sequence of pYes-FLuc-nanoLuc was verified by sequencing.

### RNA isolation, Northern blotting and signal quantification

We used the FAE method described in (Shedlovskiy et al. 2017a) to isolate RNA from cells. To isolate RNA from CFE, S180, P180, and from *in vitro* translation/*Renilla* luciferase reactions, we used TRI REAGENT-LS according to the manufacturer’s recommendations. To isolate RNA from gradient fractions, each fraction was treated with 100 μg/ml proteinase K in the presence of 1% SDS and 10 mM EDTA for 20 min at 42°C, followed by phenol/chloroform extraction and isopropanol precipitation. All RNA pellets were resuspended in FAE solution (Formamide, 10 mM EDTA) for 15 min at 65°C, shaking. RNA was separated on 1.2% formaldehyde-containing agarose gel as described in (Mansour and Pestov 2013). Prior to transfer onto Nylon membrane (GE Healthcare, cat# NS0921), gels were stained with SYBR Gold and scanned using a Typhoon 9200 imager (GE Healthcare) at 532 nm to visualize RNA. For hybridizations, we used a [^32^P]-labeled probe specific for the gene encoding TAP (5’-GCCGAATTCTCCCTGAAAA-3’), a [^32^P]-labeled probe y540 against 25S rRNA (5’-TCCTACCTGATTTGAGGTCAAAC-3’) or [^32^P]-labeled probe y500 against 18S rRNA (5’-AGAATTTCACCTCTGACAATTG-3’). We used Typhoon 9200 in the phosphorimaging mode to detect radioactive signal, which was analyzed with ImageQuant software (GE Healthcare). For quantification, the volume of the hybridization signal corresponding to the RNA specie of interest was converted to phosphorimaging units, and the background (average image background) was subtracted.

### Preparation of translationally active lysate CFE (cell-free extract)

*Saccharomyces cerevisiae* BY4741 strain was grown in 1 L of YPD medium to OD_600_∼0.8, cells were harvested by centrifugation, washed twice in H_2_O and twice in freshly prepared buffer A (20 mM Hepes-KOH (pH 7.4), 100 mM KOAc, 2 mM Mg(OAc)_2_, 2 mM DTT). The weight of the cell pellet was measured, and the pellet was resuspended in buffer A containing 8% mannitol in a 2:3 volume/cell weight ratio. Cell slurry was dripped directly into liquid nitrogen to form small ice beads. Frozen yeast/buffer beads were transferred into the prechilled grinding vial containing a metal rod and placed into a SPEX freezer mill chamber filled with liquid nitrogen. Yeast/buffer beads were powdered using the following setting: 1 min of grind, 1 min off, 8 cycles in total. The powdered cells were transferred to the prechilled 10.4 mL ultracentrifuge tube (Backman), allowed to thaw on ice, and yeast suspension was centrifuged in Beckman ultracentrifuge for 15 min at 4°C at 30,000 x g using fixed angle Beckman rotor Type 80 Ti. The clear phase between the pellet and cloudy upper lipid layer was collected (∼6 mL) and centrifuged again for 35 min at 4°C at 100,000 x g in Beckman rotor Type 80 Ti. Once again, the clear phase between the pellet and cloudy upper layer was collected, and 2.5 mL was applied on gel filtration column PD10 Sephadex G-25 (GE Healthcare) pre-equilibrated in buffer A containing 20% glycerol. For elution, we used 5 mL of buffer A containing 20% glycerol. We collected 10 fractions (500 μl each) of yeast extract. RNA content in each fraction was measured spectrophotometrically, and fractions with at least 60-75% of the highest RNA concentration were pooled, aliquoted into Eppendorf tubes (100 µL aliquots) and frozen in liquid nitrogen for storage at -80°C.

### PCR and RNA reporter preparation

PCR reactions and reporter RNA synthesis were performed as described in (Trainor et al. 2021) with minor modifications. Briefly, a sequence corresponding to a reporter gene cloned in pYes2 as described in the “ Plasmids” section was amplified with DreamTaq polymerase using forward (5’-CGGATCGGACTACTAGCAGCTG-3’) and reverse (5’-TTCATTAATGCAGGGCCGCAAATT-3’) primers that anneal upstream and downstream of the T7 promoter and the CYC1 terminator elements on pYes2, respectively. PCR products were concentrated using the ZYMO column. m7G-capped mRNA was generated using 1 μg of PCR-generated DNA template and mMESSAGE mMACHINE™ T7 transcription kit, according to the manufacturer’s recommendations. The reaction was carried out at 37 °C for 2 h, followed by DNase treatment for 15 min at 37 °C and RNA precipitation with LiCl. RNA pellet was washed with 80% EtOH, air-dried, and resuspended in 30 μL of H_2_O. Concentration was determined; RNA was aliquoted and stored at -80°C.

### Translation reactions using complete CFE

For one reaction (15 μL) we used 7.5 μL of yeast extract (final protein concentration ∼25-30 mg/mL), 1.5 μL of RNA (200-400 ng), and 6 μL of 2.5 × master mix (50 mM Hepes–KOH (pH 7.6), 25 μM each amino acid, 5 mM Mg(OAc)_2_, 125 mM KOAc, 50 mM creatine phosphate, 0.15 U creatine kinase, 5 mM DTT, 2.5 mM ATP, 0.25 mM GTP, 1 U RiboLock). Reactions were incubated at 21°C for 30-180 min (indicated in Figure legends).

### Translation reactions using RSR format

CFE was centrifuged at 180,000 × g for 2 h in the TLA55 rotor (55,000 rpm) at 4**°**C. The supernatant was collected, placed into a new tube, and stored on ice (S180). Pellet was washed with translation reaction buffer A (20 mM Hepes–KOH, pH 7.4; 100 mM KOAc; 2 mM Mg(OAc)_2_; and 2 mM DTT) (Wu and Sachs 2014), resuspended in 100 μL of the same buffer by agitation at 21°C for 30 min (or as indicated in Figure legends); centrifuged at tabletop centrifuge for 15 min at 4**°**C and ribosome suspension was transferred into a new tube. RNA concentration was measured. Alternatively, we centrifuged 100 μL of complete CFE through the 20% glycerol cushion (500 μL) prepared in translation buffer A at 180,000 × g for 2 h in TLA55 rotor (55,000 rpm) at 4**°**C. The supernatant was carefully discarded; the ribosomal pellet was processed as described above for CFE. To isolate ribosomes from cells, cell cultures were collected by centrifugation at 3,300 rpm for 3 min at 4**°**C (using Eppendorf centrifuge 5810R equipped with the A-4-62 rotor) and washed twice with buffer A. Cell pellet was resuspended in buffer A supplemented with 200 μg/mL of heparin and lysed by 10–12 cycles of 30 sec vortexing followed by 30 sec incubation on ice in the presence of glass beads (Sigma). Cell lysates were collected by centrifugation at 21,000 × g for 15 min in a tabletop centrifuge at 4**°**C; the supernatant was layered onto 20% glycerol cushion (500 μL) prepared in translation buffer A and centrifuged at 180,000 × g for 2 h in TLA55 rotor (55,000 rpm) at 4**°**C. Supernatant was discarded; the ribosomal pellet was processed as described above for CFE.

For RSR reaction with mRNA reporter, we used 6 μL of ribosome-free supernatant (S180), 1.5 μL of ribosomes (concentrations are indicated in Figure legends and in text), 1.5 μL of capped-mRNA reporter (200-400 ng) and 6 μL of 2.5 × master mix (50 mM Hepes–KOH (pH 7.6), 25 μM each amino acid, 5 mM Mg(OAc)_2_, 125 mM KOAc, 50 mM creatine phosphate, 0.15 U creatine kinase, 5 mM DTT, 2.5 mM ATP, 0.25 mM GTP, 1 U RiboLock). Reactions were incubated at 21°C for 30-180 min (indicated in Figure legends).

For RSR reaction with endogenous transcripts present in CFE, we used 6 μL of ribosome-free supernatant (S180), 1.5 μL of ribosomes (concentrations are indicated in Figure legends and in text), and 7.5 μL of 2 × master mix (40 mM Hepes–KOH (pH 7.6), 20 μM of essential amino acids minus Methionine and Cysteine, 4 mM Mg(OAc)_2_, 100 mM KOAc, 40 mM creatine phosphate, 0.12 U creatine kinase, 4 mM DTT, 2 mM ATP, 0.2 mM GTP, 0.8 U RiboLock and 1 mCi/mL EasyTag EXPRESS[^35^S]Met/Cys Protein Labeling mix). Reactions were incubated at 21°C. At 5, 30 and 45 min, 4 μL aliquots were taken from the reaction tube and added to 96 μL of 1M NaOH; tubes were incubated at 37°C for 10 min (to hydrolyze RNA) and mixed with 900 μL of ice-cold 25% TCA supplemented with 2% casamino acids to precipitate the translation product. Precipitation mixtures were applied on Whatman® GF/A glass fiber filters, washed six times with 5% TCA, once with acetone, air-dried, and placed in a scintillation vial. 2 mL of scintillation fluid were added into each vial and incorporation of [^35^S-Met/Cys] into polypeptides was determined by counting in a scintillation counter.

### Luciferase assays and statistical analysis

We used the *Renilla* Luciferase Assay System from Promega (cat#E2810) and Nano-Glo® Dual-Luciferase® Reporter Assay System (cat#N1610) according to the manufacturer’s protocol. The luminescent signal was measured on a GLOMAX 20/20 luminometer. Statistical analysis was performed by one-way ANOVA with GraphPad PRISM 5.

For RNA and protein extraction from the luciferase reaction, 100 μL of TRI REAGENT-LS reagent were added to 100 mL of the luciferase reaction. Samples were stored at -80°C prior to processing according to the manufacturer’s recommendations.

### Western blotting and signal quantification

To analyze *in vitro* translation reaction products, proteins were isolated from 15 μL of the translation reactions using TRI REAGENT-LS according to the manufacturer’s recommendations. Protein pellets were analyzed as described in (Trainor et al. 2021). In brief, protein pellets were resuspended in 50 μl of 1 x SDS-PAGE loading dye, 10 μL were resolved by SDS-PAGE using 10% polyacrylamide gels, transferred onto nitrocellulose membrane, and blocked with 10% milk in PBS–0.1% Tween 20 (PBST). Membranes were incubated with PAP antibodies or anti-Rpl3 primary antibodies, washed with PBST, and for Rpl3 blots, incubated with anti-mouse secondary antibodies. For detection, we used Crescendo chemiluminescent HRP detection reagent.

### Sucrose gradient centrifugation analysis

To analyze CFE, cellular lysate, CFE-derived or cell-derived solubilized ribosomes by sucrose gradient centrifugation analysis, 150 μg of total RNA were loaded onto 15– 45% (w/v) sucrose gradients prepared in 70 mM NH_4_Cl, 4 mM MgCl_2_ and 10 mM Tris–HCl (pH 7.4). The volume of each gradient was 11 mL. Gradients were centrifuged at 188,000 × g at 4°C for 4 h in Beckman SW41Ti rotor (36,000 rpm) and fractionated into 12 fractions (1ml each) using a Beckman fraction recovery system connected to an EM-1 UV monitor (Bio-Rad).

## ACKNOWLEDMENTS

We are grateful to Anton A. Komar, Russ Sapio, Daniel Smethurst for discussion, comments on the manuscript, and technical assistance. We thank Mike Henry for the pBS1761 plasmid, Jonathan Dinman for the pJD375 plasmid. We would like to express our gratitude to Tatyana Klimova for editing this manuscript. This study was supported by the National Institutes of Health (NIH) Grant R01GM114308 (to N. S.).

## Author contribution

D.G.P., N.S., and B.M.T. conceptualized the idea; N.S. and B.M.T. perform the experiments; D.G.P., N.S., and B.M.T. analyzed the data; N.S. wrote the original draft; B.M.T. and N.S. reviewed and edited the manuscript; N.S. provided supervision and the funding.

